# Increased genetic marker density reveals high levels of admixture between red deer and introduced Japanese sika in Kintyre, Scotland

**DOI:** 10.1101/588327

**Authors:** S. Eryn McFarlane, Darren C. Hunter, Helen V. Senn, Stephanie L. Smith, Rebecca Holland, Jisca Huisman, Josephine M. Pemberton

## Abstract

Hybridization is a natural process at species range boundaries, but increasing numbers of species are hybridizing due to direct or indirect human activities. In such cases of anthropogenic hybridization, subsequent introgression can threaten the survival of native species. To date many such systems have been studied with too few genetic markers to assess the level of threat resulting from advanced backcrossing. Here we use 44,999 single nucleotide polymorphisms and the ADMIXTURE program to study two areas of Scotland where a panel of 22 diagnostic microsatellites previously identified introgression between native red deer (*Cervus elaphus*) and introduced Japanese sika (*Cervus nippon*). In Kintyre we reclassify 26% of deer from the pure species categories to the hybrid category whereas in the NW Highlands we only reclassify 2%. As expected, the reclassified individuals are mostly advanced backcrosses. We also investigate the ability of marker panels selected on different posterior allele frequency criteria to find hybrids assigned by the full marker set, and show that in our data, ancestry informative markers (i.e. those that are highly differentiated between the species, but not fixed) are better than diagnostic markers (those markers that are fixed between the species) because they are more evenly distributed in the genome. Diagnostic loci are concentrated on the X chromosome to the detriment of autosomal coverage.

## Introduction

Hybridization is of rising concern to conservation biologists (Rhymer & Simberloff 1996; Allendorf et al. 2001; Grabenstein & Taylor 2018). While hybridization regularly occurs naturally (Mallet 2005) and is an integral part of the evolutionary history of many species (Payseur and Rieseberg 2016), anthropogenic hybridization resulting from human actions is expected to increase in frequency and intensity in the future, as introductions and range shifts due to climate change increase the occurrence of secondary contact among previously allopatric species (Crispo et al. 2011). A recent review of anthropogenic hybridization found that most cases of admixture result in the breakdown of reproductive isolation between previously distinct species (Grabenstein & Taylor 2018). Biologists have flagged anthropogenic hybridization as a conservation concern, and the rapidity with which hybrid swarms are formed (Grabenstein & Taylor 2018) suggests that this is a problem that is both common and urgent. Pure species may no longer exist as such, and the evolutionary trajectory of a species may be radically changed due to the introgression of large numbers of alleles which have already proved viable in an allopatric species.

Backcrossing of hybrid individuals with parental species leads to many admixed individuals that are difficult to differentiate from parental species individuals (McFarlane & Pemberton 2019). When hybrids mate with parental species individuals (instead of other hybrid individuals), bimodal hybrid zones can be formed, where there are many individuals with a small proportion of introgression from another species. This leads to a large proportion of individuals of hybrid ancestry in the population that are difficult to distinguish either genetically or phenotypically from parental species. In the case of genetic analysis, this is because with each generation of backcrossing a doubling of markers is needed to detect introgression with the same degree of confidence (Boecklen & Howard 1997). However, many current studies of anthropogenic hybridization use an average of 20 markers to detect introgression (Todesco et al. 2016), which is too few to reliably detect even second-generation backcrosses in most systems (Boecklen & Howard 1997; Vähä & Primmer 2006). The limited detection power of the marker panels that are typically used in studies of anthropogenic hybridization are insufficient to detect on-going introgression and it seems likely that many thousands of markers will be necessary to accurately identify the majority of hybrid individuals (McFarlane & Pemberton 2019).

The use of genomic markers (i.e. thousands of SNPs as compared to tens of microsatellite markers) increases the detection probability of admixed individuals, particularly in systems where backcrossing has occurred for multiple generations. Theoretically with each generation of backcrossing, the proportion of individuals that are homozygous at diagnostic markers for alleles from one parental species increases, with the power to detect backcrosses sharply decreasing with each additional generation (Boecklen & Howard 1997). For these reasons it has been suggested that when determining how many markers to genotype researchers consider not only divergence between hybridizing species (as suggested by (Vähä & Primmer 2006)), but also the number of elapsed generations since hybridization has begun and the recombination rate of the taxa studied (McFarlane & Pemberton 2019). Empirically, use of high-density markers has indeed led to an increase in detection of backcrossed individuals. For example in Italy where wolves and dogs have hybridized, 1 – 5% of wolves were found to be hybrid individuals when 16 – 18 microsatellite markers were used (Randi 2008), whereas 62% of sampled individuals were determined to be hybrid when 170 000 SNP markers were used (Pilot et al. 2018). Although the individuals sampled were not the same, and the study goals differed, it is telling that there is such a discrepancy in the estimated proportion of hybrid individuals when a high density marker panel was used. Similarly, all individuals in a westslope cutthroat trout – rainbow trout hybrid zone that were previously thought to be parental species were discovered to be hybrid individuals when re-analyzed with more markers (Boyer et al. 2008; Hohenlohe et al. 2013). Use of higher density marker panels has the potential to change the qualitative and quantitative understanding of hybrid systems, including providing information on the direction of backcrossing, the number of generations since backcrossing began and the distribution (e.g. hybrid swarm, unimodal or bimodal) of the hybrid system (McFarlane & Pemberton 2019).

Even with the use of genomic markers, not all markers will contribute equally to hybrid detection. Diagnostic markers (DM), those that have completely fixed allele differences between populations, have more power than markers that are polymorphic in either population (Pritchard et al. 2000; Vähä & Primmer 2006). Ancestry informative markers (AIM), which have substantial allele frequency differences between parental populations are also more powerful than markers which are highly polymorphic within one or both parental populations (Shriver et al. 2003). Thus, if a situation requires the use of fewer markers (e.g. to lower the cost of genotyping individuals, or because of the need for rapid testing (Senn et al. 2018)), it would be best to focus on diagnostic or ancestry informative markers, with the caveat that more markers will always lead to improved detection of hybrid individuals (McFarlane & Pemberton 2019). For example, Galaverni et al. (2017) compared 48, 96, and 192 AIMs taken from a set of approximately 25,000 markers that had been thinned for linkage disequilibrium (LD). They found that each of these marker sets performed similarly, although there were no DM included in this comparison. A comparison of hybrid detection that includes both diagnostic markers and ancestry informative markers would contribute to confidence when researchers use AIM, especially since there are not always DM available.

It should be noted that both diagnostic and ancestry informative markers can have a biased distribution across the genome due to the evolutionary history of a population. If there is heterogeneity in introgression across the genome, then using too few markers may miss hot-spots of introgression. For example, genes that are associated with post zygotic reproductive isolation are expected to be concentrated on sex chromosomes, due to the smaller effective population size and limited recombination of sex chromosomes (Qvarnström & Bailey 2009). Empirically, a disproportionate number of genes associated with reproductive isolation have been traced to the sex chromosomes (Coyne & Orr 2004), including in mouse hybrids (Payseur et al. 2004), swordtail hybrids (Schartl 1988), and flycatcher hybrids (Sæther et al. 2007). Markers in strong LD with such genes are likely to be diagnostic (DM). When selecting markers for hybrid detection, the exclusive use of diagnostic markers will ignore introgression in other parts of the genome, where DM are not present. Further, thinning for LD is likely to exclude many DM, decreasing the power of the analysis to detect hybrid individuals. AIM may also be disproportionately distributed across the genome, but perhaps not with the same intensity as markers that have fixed between species.

Hybridization between native red deer (*Cervus elaphus*) and Japanese sika (*C. nippon*) in Scotland is a scenario in which the use of high-density markers to explore the extent of backcrossing seems very likely to provide new insights. Sika were introduced from Japan to Ireland in 1860 and from Ireland to Scotland beginning in 1870 (Ratcliffe 1987). As the hybridization that occurred after these introductions was facilitated by human actions, this is considered anthropogenic hybridization (Grubenstein and Taylor 2018, Allendorf 2001). Based on earlier work using microsatellite markers, there appear to be two different scenarios of admixture between red deer and sika in Scotland, which we have studied here using two separate populations. Sika were introduced to the Kintyre Peninsula in 1893, and phenotypically hybrid individuals were first reported in the region in the 1974 (Ratcliffe 1987). Given a generation time of approximately 5 years in red deer (Coulson et al. 1998), we might expect at least 6 -7 generations of backcrossing in this system, as our dataset includes deer sampled as early as 2006. However, we cannot discount the possibility of hybridization prior to introduction to Scotland, as hybridization was recorded in the Irish source population in the 1880’s (Powerscourt 1884). Previous research in Kintyre used 22 diagnostic microsatellites to determine that there is a range of admixture proportions and that backcrossed individuals are common (Senn et al. 2010a; Smith et al. 2018). The use of a diagnostic mitochondrial marker demonstrated the existence of backcrosses with very low levels of nuclear introgression, beyond the detection of the microsatellite markers (Smith et al. 2018). Hybrid individuals have phenotypes that are closely correlated with their admixture score (Senn et al. 2010b). While intermediate hybrids may be detectable by phenotype, advanced backcrosses are easily confounded with the parental species, and are difficult for deer hunters (“stalkers”) to identify in the field (Senn & Pemberton 2009; Smith et al. 2014).

Sika were introduced to the north of Scotland between 1889 and 1900 (Ratcliffe 1987). Hybrid individuals have been documented in this region, but in contrast to Kintyre, there does not seem to be a hybrid swarm - the few hybrids detected by microsatellites and mtDNA are the result of extensive backcrossing (Smith et al. 2018). In this study we screened populations in the NW highlands, where these previously-detected hybrids were sampled.

Anthropogenic hybridization between red deer and Japanese sika provides an opportunity to use SNP markers to search for cryptic backcrossed individuals that were previously identified as parental individuals. Using substantially more markers than were previously available, this study has three aims: (1) to determine how many individuals are hybrids when scored using greatly increased marker density; (2) to determine the distribution of hybridization and introgression in each study area, specifically whether the hybrid zone is uni or bi directional and whether hybrids mate with other hybrids or with parental species individuals; (3) to study the performance of different classes of marker (diagnostic, ancestry informative and random) in determining hybrid status and understand the difference arising from marker number and genomic location.

## Methods

### Study area and sampling

Samples were collected in the Kintyre peninsula, and from the northwest highland region of Scotland (Figure 1). Samples were collected from 15 forestry sites by Forestry Commission Scotland stalkers as part of routine culling operations. The Kintyre samples included 336 individuals shot in 2006-2007 and 187 shot in 2010-2011. The NW highland samples included 108 individuals sampled in 2010-2011. All samples were shot as encountered, i.e. without bias in terms of phenotype (Smith et al. 2018). Sample collection consisted of ear tissue and has previously been described in Senn and Pemberton (2009) and (Smith et al. 2018). Samples were either preserved in 95% ethanol or frozen.

**Figure 1:**
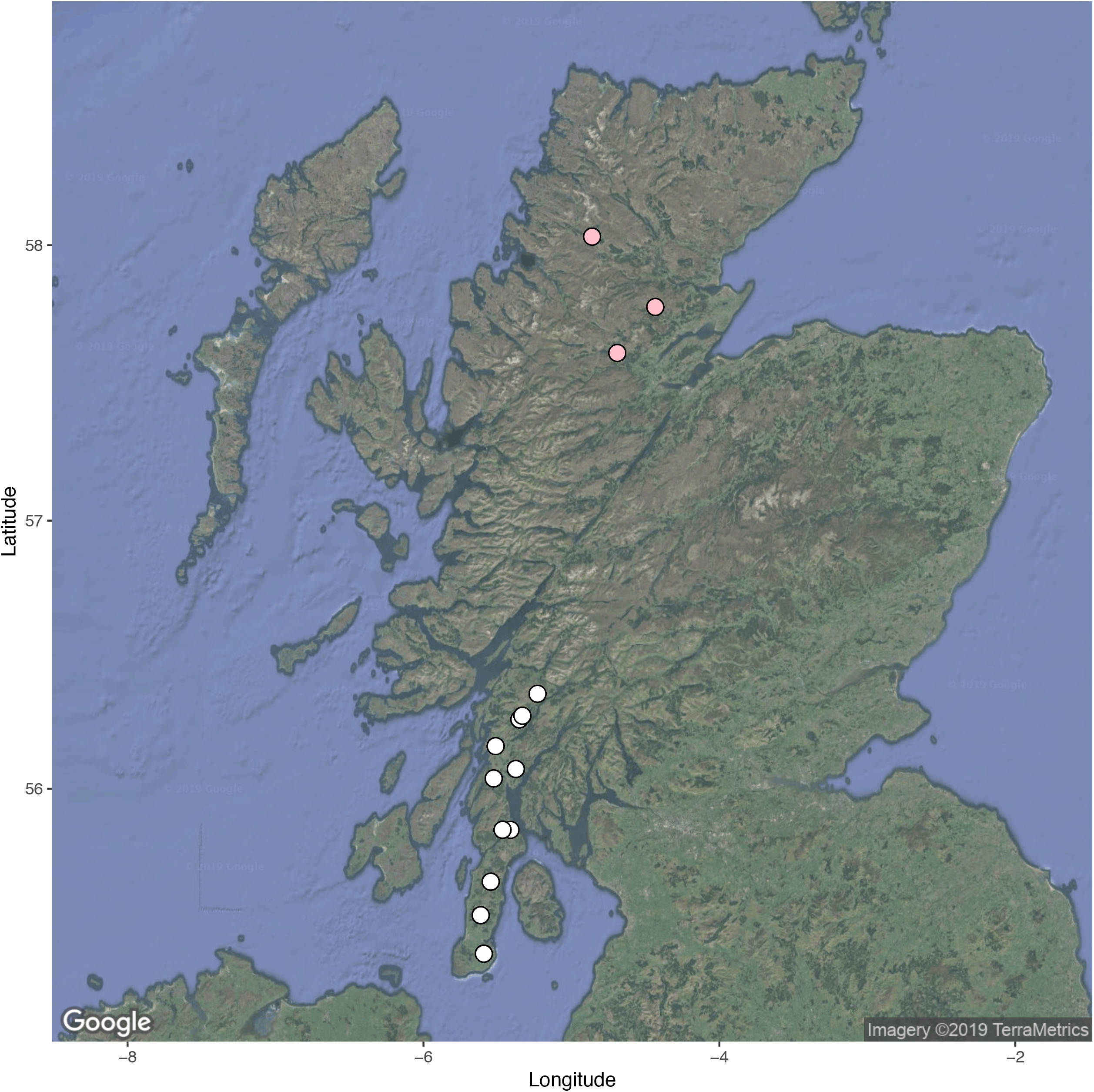
A map of Scotland, showing the approximate sampling locations for deer from Kintyre (white) and the NW Highlands (pink) that were genotyped on a 50K cervine SNP array. The map is from Google Maps, accessed using ggmap (Kahle & Wickham 2013).

### Microsatellite genotyping, mtDNA haplotyping

All individuals included in this study were previously genotyped at 22 diagnostic microsatellites (Senn et al. 2010a; Smith et al. 2018). Most individuals were also typed at the mitochondrial DNA control region, which includes a 39-bp sequence with a variable number of tandem repeats, of which red deer have a single repeat and Japanese sika have three repeats (Cook et al. 1999; Smith et al. 2018).

### DNA extraction for SNP analysis

We used the DNeasy Blood and Tissue Kit (Qiagen) according to the manufacture’s instructions to re-extract DNA for SNP analysis, except that to obtain DNA at a sufficiently high concentration, in the last step we eluted twice in 25 µl TE. DNA concentration was assayed using the Qubit™ dsDNA BR Assay Kit (Invitrogen™) and any samples below 50ng/µl were vacuum-concentrated, re-extracted or omitted from SNP analysis.

### SNP Genotyping

SNP genotyping was carried out using the cervine Illumina iSelect HD Custom BeadChip using an iScan instrument following manufacturer’s instructions (as in Huisman et al. 2016). To develop this chip five European red deer, and two wapiti (*Cervus canadensis or Cervus elaphus canadensis*) were whole genome sequenced (Brauning et al. 2015), in addition to two pools of 12 pure species red deer from the Isle of Rum, Scotland and 1 pool of 12 pure species sika (based on microsatellite data) from Kintyre, at 10x coverage. The majority of SNPs on the chip (53K attempted loci) were selected to be polymorphic within red deer, specifically within the Rum red deer. However, during the development of the chip 2250 SNPs were included that were expected to be diagnostic between red deer and sika, and 2250 SNPs were included that were expected to be diagnostic between red deer and wapiti. Within each marker set, SNPs were selected to be evenly spaced throughout the genome according to map positions in the bovine genome, with which the deer genome has high homology (Johnston et al 2017, Slate et al 2002). To check for consistency between batches and calculate the repeatability of each SNP, we used the same positive control twice on each 96-well plate (Huisman et al. 2016). Genotypes were scored using GenomeStudio (Illumina), using the clusters from (Huisman et al. 2016). SNPs that didn’t successfully cluster according to Huisman (et al. 2016) were clustered manually. Quality control was carried out in PLINK (Purcell et al. 2007). We excluded individual samples with a call rate of less than 0.90. We deleted loci with minor allele frequency less than 0.001, and/or call rate of less than 0.90. We did not exclude SNPs based on the Hardy-Weinberg equilibrium, as any diagnostic markers fixed between the two species are not expected to be in HWE in an admixed population.

### Estimating global F_ST_

To estimate global F_st_ between red deer and sika in our two study areas, we considered only individuals that had been assessed as parental species from the phenotype by the ranger concerned. We used phenotype rather than the genotype from our ADMIXTURE analyses (see below) to exclude admixed individuals for this analysis to avoid a circular argument regarding the power we had to detect hybrid individuals. We used the R package ‘assigner’ (Gosselin et al. 2016) to estimate Weir and Cockerham’s F_st_ (Weir & Cockerham 1984), with 95% confidence intervals (CI) based on bootstrapping.

### Estimating admixture

For the microsatellite data, we used the STRUCTURE Q scores from Smith et al. 2018 to compare species assignment with the Q scores estimated using SNP markers (described below). However, unlike Smith et al. (2018), we used credibility intervals rather than a threshold cut off to assign individuals to species categories (see description below).

For the SNP data, we used ADMIXTURE (Alexander & Lange 2011) to estimate the admixture (Q) score for each individual. ADMIXTURE uses maximum likelihood to estimate individual ancestry proportions (Alexander et al. 2009). We ran ADMIXTURE unsupervised separately on the deer from Kintyre and those from the NW highlands, as these are two separate populations and we did not want to inadvertently introduce population structure while examining hybridization (Gompert & Buerkle 2016). We assumed two ancestral populations. We used the bootstrapping function in ADMIXTURE to estimate standard errors, from which we calculated 95% confidence intervals as 1.96*SE. We also used fastSTRUCTURE (Raj et al. 2014) to estimate Q scores which were highly correlated with the Q scores from ADMIXTURE (Pearson’s correlation coefficient R= 0.9999, t=6720.1, df=631, p<2.2e-16), but used ADMIXTURE Q scores in all the analyses reported below in order to report error around point estimates (McFarlane & Pemberton 2019). It should be noted that neither analysis method accounts for linkage between markers. When we excluded markers based on LD (r2<0.3, 12K SNPs remaining), we found a very high correlation between admixture scores with and without this LD pruning (using ADMIXTURE, Pearson’s correlation = 0.9915, t=172.1, df=511, p<2.2e-16), and that only 6 individuals were assigned to a difference species in the LD pruned analysis. Thus, all downstream analyses were done without accounting for LD.

To categorize animals into pure red deer, pure sika deer or hybrid, we determined whether the 95% confidence intervals for the Q scores overlapped 0.99999 for pure red deer, or 0.00001 for pure sika deer. These are the highest and lowest possible Q scores produced by ADMIXTURE, and, with many markers, confidence intervals were extremely narrow. All other individuals were classified as ‘hybrids’. These stringent criteria for assigning an individual to a parental species means that we are more likely to wrongly assign parental species individuals as hybrids than wrongly assign hybrid individuals as parental (i.e. low accuracy or a Type II error with regard to wrongly assigning individuals as parental species vs. low efficiency or a Type I error as defined by (Vähä & Primmer 2006)).

### Effect of varying SNP selection

We wanted to compare the efficiency of using our entire marker panel against using either only diagnostic markers, those fixed for different alleles in each parental population, ancestry informative markers, those with distinct allele frequency differences in the two parental populations (Shriver et al. 2003), or markers indiscriminately sampled from the SNPchip that were neither diagnostic nor ancestry informative (hereafter ‘random’). Diagnostic markers (DM) were defined as having posterior allele frequencies (in the ADMIXTURE analysis) of more than or equal to 0.99999 in either direction. Ancestry informative markers (AIM) were defined as having extreme posterior allele frequencies in each population, estimated as |p1-p2|>0.95, where p1 is the allele frequency in one population, and p2 is the allele frequency in the other population.

To compare the power of each marker type, without confounding by the number of markers used, we first determined the number of DM that were available, which was 629, and then randomly sampled this number of markers from each of the AIM (excluding DM) and the entire marker panel (excluding DM and AIM). We also included a marker set that was ten times as many randomly sampled SNPs (6290) as well as the entire SNP panel (approximately 45 thousand SNPs) and the 22 microsatellite markers used in Smith et al. 2018. We refer to these datasets as 45K, 629 DM, 629 AIM, 629 random, 6290 random later in the text. We then reran the ADMIXTURE analysis with each of the five marker sets for the Kintyre sample and re-estimated the number of individuals in each class (red, sika, hybrid). We focused on the deer from Kintyre because the sample size was larger and there was prior evidence of a large number of hybrid individuals in this region (Senn et al. 2010a), a finding confirmed below. To understand the uncertainty around the number of individuals in each class generated by the different marker sets, we ran each analysis 100 times, and varied the ‘set seed’ in ADMIXTURE (-s time) to allow for slight differences between replicates. Note that the 629 AIM and the 629 random and 6290 random panels were a new random draw of markers for each bootstrap in the analysis. We present the mean estimated number of red deer, hybrid and sika called, as well as the standard errors around each estimate from the replicated analyses.

For each category of marker, we compared the number of deer per iteration that were identified to species class differently when compared to the 45K marker set. We refer to these deer as ‘mismatches’. We used a linear model with the number of mismatches as the response variable explained by the marker category to ask which marker category gave results most similar to those from the 45K panel. We report the mean number of mismatches ± standard error, as well as an ANOVA of the linear model. Since we found large differences between marker groups, we explored the location of markers using the red deer linkage map (Johnston et al. 2017) as a possible explanation of the observed differences.

All data and code are available at https://figshare.com/projects/Increased_genetic_marker_density_reveals_high_levels_of_admixture_between_red_deer_and_introduced_sika_in_Kintyre_Scotland/66860.

## Results

After quality control, there were 513 individual deer from ten sampling areas on the Kintyre peninsula, 108 deer from three sites in the NW Highlands, and 44 999 SNPs (Figure 1). After excluding individuals that were assessed to be hybrids by rangers (N = 30 on Kintyre and N= 27 in the NW highlands), and using all markers, F_st_ between the two species was 0.5321 (95% CI = 0.529 – 0.534) in Kintyre and 0.4751 (95% CI = 0.472 – 0.478) in the NW Highlands. Note that the inclusion of putatively diagnostic markers is likely to have inflated F_st_ compared with the true value.

### Kintyre

The use of SNP genotyping led to a different classification of many animals when compared to microsatellite genotyping. Using the definitions outlined above, we found 159 pure red deer (i.e. those with a CI that overlapped 0.99999), with an average Q score of 0.9986+/−0.003 (+/− SD), 132 pure sika deer (CI overlapped 0.00001), with an average Q score of 0.00169+/− 0.003 and 222 hybrid deer with an average Q score of 0.6144+/− 0.393 (Figure 2a). Hybrid individuals had Q scores that spanned from 0.008 to 0.992, consistent with 6-7 generations of backcrossing. In comparison to previous work that used 22 microsatellites (e.g. Smith et al. 2018), we classified many more individuals as hybrids (N= 222, vs 119 with microsatellites), and the majority of newly classified hybrids had a Q score of >0.85 (red deer like hybrids) or <0.15 (sika like hybrids). This is in spite of a high correlation of Q score estimates between microsatellites and SNPs (R^2^= 0.997, p<2.2e-16). Additionally, when including Q score as a covariate in a linear regression, we found that newly detected hybrids have a more southerly distribution than previously detected hybrids (ANOVA, F_1, 221_=4.70, p=0.031). This pattern of many individuals with low levels of introgression is consistent with a bimodal hybrid zone, as has been demonstrated in previous studies (Goodman et al. 1999; Senn & Pemberton 2009). We found three potential F1 individuals, for which 95% CIs overlapped Q=0.5, but on further examination of the heterozygosity of diagnostic markers we determined that these individuals were advanced hybrids with intermediate Q scores, since these individuals were not heterozygous for a high proportion of diagnostic markers. Finally, of the hybrid individuals that we identified, 167 had a red deer mitochondrial marker, and 45 had a sika mitochondrial marker (mtDNA was not available for 12 individuals). This is in contrast to 103 hybrids with red mtDNA and 13 hybrids with sika mtDNA markers found using microsatellite markers (89.1% red mtDNA, 10.9.% sika mtDNA with microsatellite data vs. 78.7% red mtDNA and 22.2% with SNP data; Pearson’s chi squared=4.51, df=1, p=0.034). Additionally, we found nine individuals that we assessed using ADMIXTURE as pure species sika, but which had red deer mitochondrial markers, indicated that some advanced backcrosses were beyond the detection limit of the full SNP panel.

**Figure 2:**
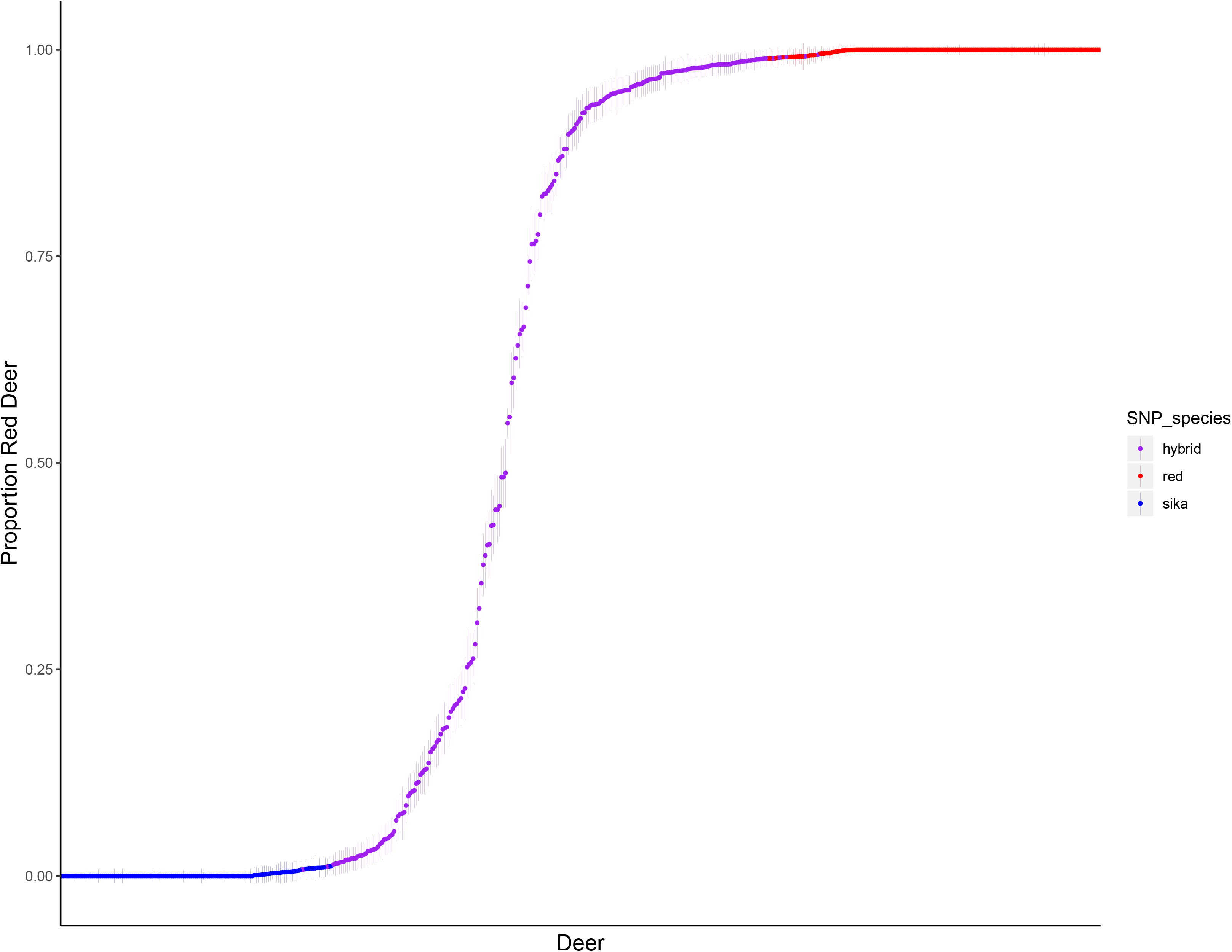
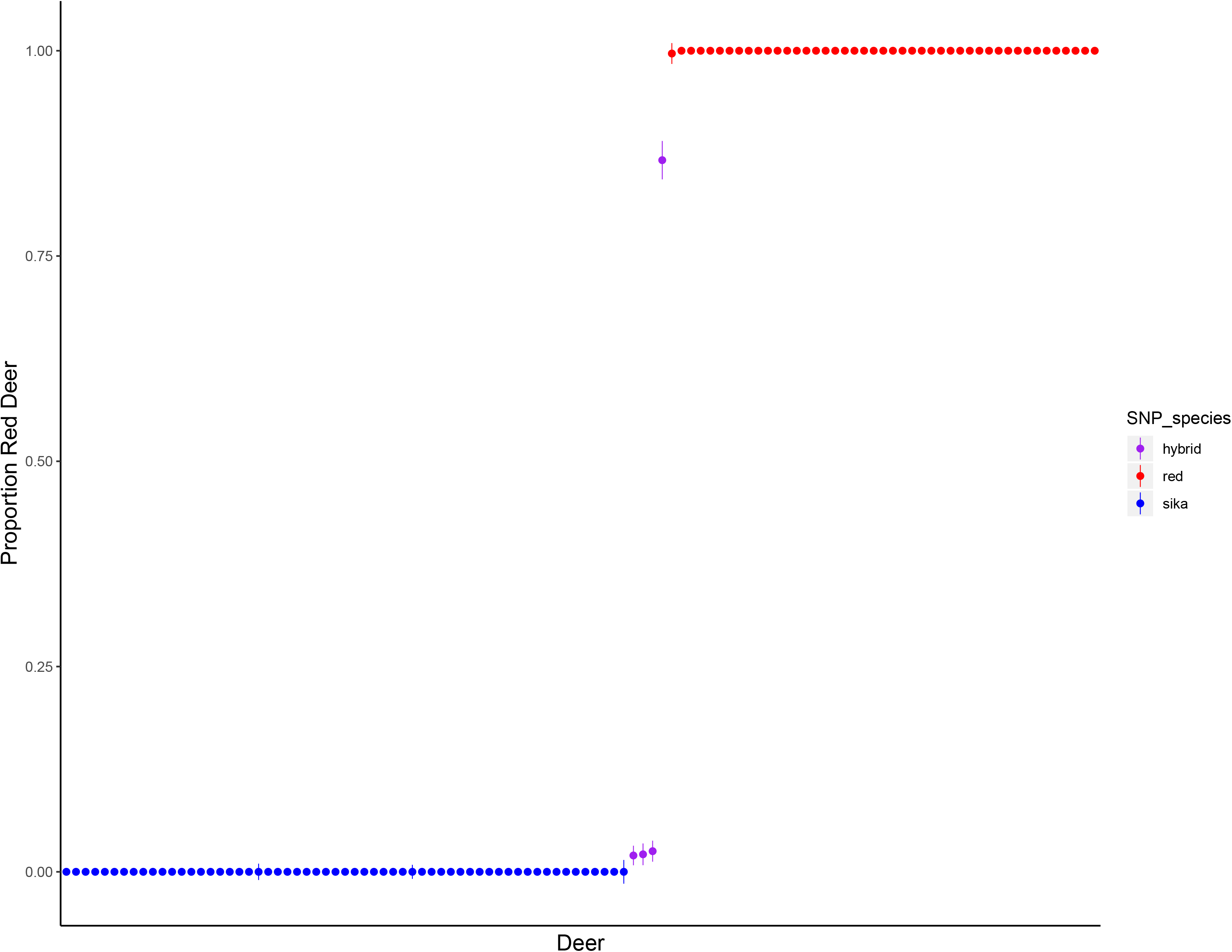
Estimates of admixture proportion (Q scores) and 95% confidence intervals for individual deer from (A) Kintyre and (B) the NW Highlands using 44,999 SNPs. Individuals are arranged by Q score following ADMIXTURE analysis. Individuals were assessed as members of the parental species if the 95% CI overlapped either 0.99999 (red deer; shown in red) or 0.00001 (sika, shown in blue), otherwise individuals were assessed as hybrid (purple). Horizontal grey lines indicate 0, 0.05, 0.95 and 1, where 0.05 and 0.95 were the thresholds used to assign species in Smith et al. 2018.

### NW Highlands

In the NW Highlands, we found 45 red deer (mean Q=0.9999+/−0001), 59 sika (mean Q=0.00001+/−0.0000), and 4 hybrids (Q=0.2333+/−0.4223). This is similar to the 45 red deer, 61 sika and 2 hybrids that were previously reported in this group of individuals (Figure 2b). The Q scores for three of these hybrid individuals ranged from 0.0199±0.006 to 0.0251±0.007, i.e. they were very sika like hybrids, while the 4^th^ individual had a Q score of 0.8667±0.133 (red like hybrid). Two of these hybrid individuals had red deer mtDNA; the other two had sika mtDNA. We also found 4 individuals that we assessed as pure species sika using the nuclear genome but that had red deer mitochondrial markers. Given the scarcity of hybrids in this region, we did not conduct further analysis on this dataset.

### Diagnostic and Ancestry Informative Marker Panels in Kintyre

We found substantial differences in the number of individuals assigned to each species by each of five marker panels (all 45K, 629 DM, 629 AIM, 629 random sample, 6290 random sample). Generally, we found that the 629 diagnostic markers and the sample of 629 random markers estimated fewer hybrids in the population than the other marker groups (Figure 3). We found that the number of mismatches (deer identified differently by one of the marker subsets than by the full 45K panel) was affected by marker type such that the 629 AIM had the fewest mismatched individuals (23.91±0.30), followed by the 629 DM (71.89±0.32), the 6290 random sample (82.27±9.0) and 629 random subsample of markers (95.55±5.8). The marker type used explained 20.4% of the variance in mismatches (F_3,396_=33.8, p=2.2e-16).

**Figure 3:**
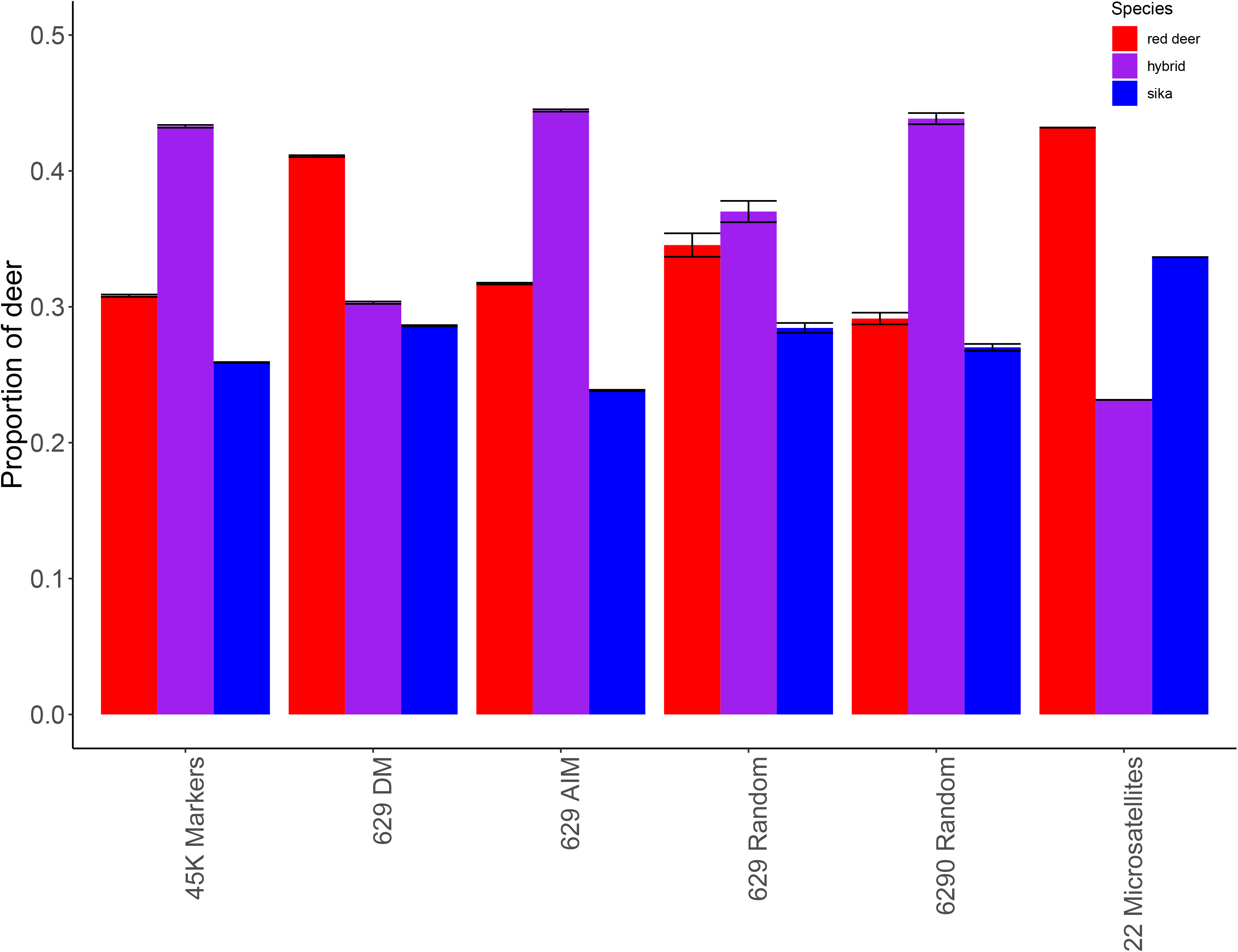
The number of deer assigned to each species category (red deer, hybrid or sika) according to all 44,999 (45K) markers, a panel of 629 diagnostic makers (629 DM), panels of 629 ancestry informative markers (629 AIM), or sets of either 629 or 6290 random markers (629 and 6290 random). These marker panels are then compared to the microsatellite results for the same individuals from Smith et al. (2018). Markers were assessed as either AIM or DM using posterior allele frequencies in ADMIXTURE (see text). Each assessment illustrated was run 100 times to allow for variation in the chosen panels (except for the DM panel) which numbered 629 markers, and with different start points for each analysis. Markers are non- nested, such that AIM do not include DM, and the random subsets of markers do not include DM or AIM. Red deer are depicted in red, hybrids are purple and sika are blue.

Inspection of the genomic location of the DM and AIM marker groups revealed a large contrast in genomic location. In the DM group, 269 of the 629 markers (42.8%) were on the X chromosome, while the autosomes had uneven coverage and chromosomes 12, 16, 22, 23 and 25 had no markers at all. In the AIM group, 186 of the 3205 markers (5.8%) were on the X and all chromosomes were marked, though of course this may not have been true of the individual draws in the 629 AIM analyses. Among all markers passing QC, 2360 of 44,999 (5.2%) were on the X. Among those markers originally selected to be diagnostic for sika and actually diagnostic for sika, a remarkable 152/282 or 53.9% were on the X.

## Discussion

We found many more hybrid individuals on Kintyre using 45K SNPs than had previously been reported using 22 microsatellites. The majority of the newly discovered hybrids were those with either a very high or very low Q score, suggesting that we mainly have higher efficiency (i.e. we are more likely to assign true hybrids as hybrid individuals; Vähä & Primmer 2006) than previous studies of this system (Senn et al. 2010a, Smith et al. 2018). However, it should be noted that we estimated a very similar F_st_ as that reported in previous studies (0.53 compared to 0.58, using microsatellite markers selected to be diagnostic (Senn & Pemberton 2009), suggesting that while increasing the number of markers allowed us to detect more hybrid individuals, more markers did not greatly affect the estimate of F_st_.

A possible concern is that AIMs do not represent hybridization, but rather incomplete lineage sorting (Sang & Zhong 2000), and as a result we are identifying some pure individuals as hybrids. However, as the use of AIMs alone would result in relatively greater uncertainty, and we have assigned hybrids based on credible intervals that do not overlap with 0 or 1, an individual with both an extreme point estimate and high uncertainty would be assessed as a parental species. As the uncertainty due to AIMs is thus built-in to our analyses, we believe that we have been conservative in our assignment of hybrid individuals. Furthermore, the existence of many highly introgressed individuals was expected due to the detection in Kintyre of ‘mitochondrial hybrids’ with microsatellite genotypes suggesting pure sika but with mitochondrial haplotypes of red deer (Smith et al. 2018), and from considerations about the power of the small number of microsatellites used in that study to detect hybrids (see McFarlane and Pemberton 2019). For these reasons, incomplete lineage sorting is unlikely to account for the increased number of hybrid individuals we have assigned here.

Even with 45K markers, we are unlikely to detect all of the backcrosses that could be present in the Kintyre population. The most extreme Q score that we found in a hybrid (i.e. an individual with credible intervals that excluded 0 or 1) was 0.0078±0.003, which would be the approximate average Q score for a sixth generation backcross. Given that hybridization was already apparent 6 – 7 generations ago (Ratcliffe 1987), there could be more extreme hybrid individuals that we could not detect. Indeed, there were thirteen individuals that we assessed as pure species sika, but that have red deer mitochondrial markers. These individuals appear to be backcrossed beyond our power to detect them using nuclear markers. Smith and colleagues (2018) also detected these individuals as mitochondrial hybrids. With 629 diagnostic markers, we would expect to detect 91.5% of 8^th^ generation backcrosses (McFarlane & Pemberton 2019), but this assumes unlinked markers. However, we are certainly getting additional power to detect backcrosses from the other, non-diagnostic markers, so it is difficult to be certain where our detection ability tapers off. Theoretically, individuals that are the product of consistent backcrossing since an initial F1 hybridization event could be homozygous for parental alleles from one species for all informative markers that we have used here, making them impossible to detect (Boecklen & Howard 1997).

A surprise in our analysis was that the diagnostic marker panel of 629 markers performed worse, in terms of finding hybrids, compared with equivalent-sized panels of ancestry informative markers (Figure 3). Diagnostic markers are the gold standard in the field (Pritchard et al. 2000), so this observation requires explanation. 71.89±0.32 individuals were mis-categorized by the 629 DM panel when compared to the full marker panel, and these animals had an average Q score of 0.976±0.02 for red like hybrids, and 0.018±0.01 for sika like hybrids, suggesting that the DM panel missed highly backcrossed individuals. The 629 AIM panel, despite comprising slightly less diagnostic markers probably outperformed the 629 DM panel because of a better distribution of the markers across the genome. Any introgression that occurred on a chromosome without diagnostic markers (12, 16, 22, 23 and 25), or in unmarked regions within the other chromosomes, would not be detected by the DM markers, and the individual would not be assigned as a hybrid. Markers not in the DM or AIM categories, here called ‘random markers’ generated more mismatches with the full marker set than the 629 AIM panels, and the smaller sample size of 629 such markers was worse in this regard than the larger panel of 6920 such markers. This is as expected if the markers are relatively uninformative but power is gained by having more of them.

The question arises as to why so many of the diagnostic markers were on the X chromosome. Although the selection criteria during the construction of the cervine SNP array included even spacing across the chromosomes (against the bovine map), both within the selected red deer polymorphic and the sika diagnostic loci, we suspect that the categorization into diagnostic versus ancestry informative markers has led to a concentration of X-linked markers into the diagnostic category. Since introgression is impeded on the sex chromosomes because of limited recombination, there may be long runs of fixed markers on the sex chromosomes in any hybridizing system (Barton 1979; Baack & Rieseberg 2007). While thinning markers for linkage will alleviate this problem to some extent, it will also limit the number of informative markers in an analysis, and thus might not be the best solution when more markers will lead to higher power. Many researchers in hybrid systems use anonymous and unmapped markers (e.g. from RADseq SNP discovery) and select diagnostic markers for analysis. Given the demonstrable downside to using diagnostic markers in our system (concentration on the X at the cost of autosome coverage, leading to fewer identified hybrids) we suggest that future analyses of anthropogenic hybrid zones should include both DM and AIM, and probably all possible markers, in their analyses. We anticipate that this will lead to increased discovery of hybrid individuals, as it has done in the red deer – sika bimodal hybrid zone on Kintyre.

The presence of so many previously undetected backcrossed individuals in Kintyre has conservation and evolutionary implications for red deer across Scotland. On Kintyre, we detected 103 hybrids among the 394 individuals that had previously been determined to be parental species. In contrast, we only classified 4 individuals (of 106) from the NW Highlands as hybrids using higher density markers. This suggests that reproductive isolation is still strong in the NW highlands, but has broken down to some extent in Kintyre, where a bimodal hybrid zone has formed. This does not necessarily mean that parental type sika and parental type red deer are regularly mating with each other, especially as we identified no F1 individuals in our data set. However, it is the case that there is the opportunity for substantial genetic exchange between red deer and sika in Kintyre. Further, since Kintyre is a peninsula attached to the Scottish mainland, there is the opportunity for it to become a source population from which admixed individuals can disperse. Previous work using microsatellite markers found a significantly higher number of advanced backcrossed individuals when moving away from the point of introduction and but no evidence of more hybridization or introgression over a 15-year period (Senn et al. 2010a). The newly detected hybrids tend towards a more southerly distribution than those hybrids that had previously been detected, although there were also newly detected hybrids in the north of Kintyre. These backcrossed individuals in the north and south regions of Kintyre provide an opportunity for sika gene flow into red deer in the rest of Scotland, and thus the possibility that pure red deer could eventually become extinct in mainland Scotland.

Whether such a scenario occurs depends on the detailed evolutionary dynamics of the system, which at the moment we can only speculate about. In the Scottish landscape, sika are proving extremely successful in commercial timber forests, and stalkers find them challenging to manage by shooting due to their secretive behaviour, so introgressing sika alleles could confer these traits on red deer. Introgressed individuals (as detected by microsatellites) have intermediate phenotypes in terms of appearance and size (Senn et al 2010a; 2010b), though differences in fecundity or female pregnancy rate were not detectable in the latter study. However, we are currently ignorant about how selection is operating on introgressed alleles and phenotypes. Future work could use higher density markers to re-examine the cline dynamics (both spatial and temporal) in Kintyre to determine if there are temporal changes over longer time spans than hitherto studied, for example if the tension zone of hybridization has moved over time (Barton & Hewitt 1985). In addition it should be possible to study whether particular genome regions are introgressing faster than others and potentially what traits they underpin.

Finally, it is a debatable matter whether there is conservation management value in detecting such advanced backcrosses. If an individual is less than 0.08% (i.e. a 6-7^th^ generation backcross as we have detected in the present study), should this individual be considered a hybrid when a management decision is made? This is a difficult question to answer (Allendorf et al. 2001), and we believe it should be answered based on policy and biological considerations, rather than determined based on the statistical power a given set of markers allows. A gene based theory of conservation has been proposed (Petit 2004), in which individuals of conservation concern are determined to be hybrids based on the alleles carried for specific genes of interest for the native parental population (i.e. rare alleles, or alleles associated with distinct phenotypes). However, such ‘gene targeted conservation’ is probably out of reach for this and many other study systems of conservation concern (Kardos & Shafer 2018). More practically, a mixed strategy of preserving those individuals that are genetically parental species (i.e. narrow Q score CIs that overlap with 0 or 1) *and* score highly on emblematic red deer phenotypes (e.g. summer coat without spots, long pointed ears, large antlers) could reduce the threat to red deer from sika introgression. A similar strategy is currently in use to preserve both the genetics and phenotypes of Scottish wildcats, where individuals with both high phenotypic and genetic scores are preferentially protected and used in a captive breeding program (Senn et al. 2018). Given the presence of so many advanced backcrosses in this system, any management considered should account for this introgression.

